# The Impact of Smoking Status on the Genomic Landscape of Lung Squamous Cell Carcinoma

**DOI:** 10.64898/2026.05.29.728662

**Authors:** Sahil Garg, Ravi Salgia, Ramya Muddasani, Lauren Antrim, Matthew Lee, Jyoti Malhotra, Danny Nguyen, Arya Amini, Yufei Liu, Sagus Sampath, Colby Jenkins, Adam Rock

## Abstract

**Purpose:** Comprehensive genomic profiling (CGP) has changed the treatment paradigm for non-small cell lung cancer (NSCLC) with the advent of molecularly targeted therapies for actionable genomic alterations (AGA). Despite this, the use of CGP is suboptimal, particularly in squamous cell lung cancer (sqNSCLC), which is more closely associated with smoking exposure and a lack of AGAs. We hypothesized that the prevalence of AGAs is inversely correlated with the chronicity and extent of smoking exposure in patients with sqNSCLC.

**Experimental Design:** We retrospectively evaluated all patients with liquid biopsy testing via Guardant 360CDX or Guardant360 in the context of any sqNSCLC diagnosis at the City of Hope Comprehensive Cancer Center between 10/2020 and 7/2023. The data was obtained on 2/23/24. Social and clinical histories were evaluated to assess the frequency of AGAs in patients with no or remote smoking history.

**Results:** Of the 56 patients in the initial evaluation, 24% (n=13) were non-smokers or remote smokers (greater than 20 years from cessation). Of these 13 patients, eight (61.5%) harbored AGA. Of these 8 patients, alterations observed included *EGFR* exon 19 deletion (50%, n=4), *MET* exon 14 skipping mutation (25%, n=2), *EGFR* G719S (13%, n=1), *EGFR* E114K (13%, n=1). Of those patients harboring AGAs that received NCCN-concordant matched targeted therapy, the objective response rate (ORR) with targeted agents was 50% and the clinical benefit rate (CBR) was 83.3%.

**Conclusions:** These data support the use of CGP in sqNSCLC particularly in patients with remote or no smoking exposure.

**Statement of translational relevance:** These data demonstrate high frequency of actionable genomic alterations (AGAs) in patients diagnosed with squamous cell lung cancer (sqNSCLC) with remote or no smoking history. Specifically, enrichment of *EGFR* and *MET* gene alterations were observed. These findings support the use of comprehensive molecular profiling in sqNSCLC. Furthermore, treatment outcomes demonstrate frequent objective responses and high clinical benefit rate supporting the use of targeted therapies in sqNSCLC harboring AGAs.

This analysis provided rationale for further research of larger datasets investigating therapeutic approaches in sqNSCLC, which may have significant implications for consensus guideline recommendations and routine clinical practice.

## Introduction

Lung cancer is the leading cause of cancer-related deaths in the United States(1). Broad histologic categories include small cell lung cancer (SCLC) and non-small cell lung cancer (NSCLC). NSCLC can be further sub-classified into two predominant histological categories: adenocarcinoma (LUAD) and squamous cell carcinoma (sqNSCLC). These histological subtypes exhibit distinct and shared clinical and histopathological characteristics. LUAD predominantly occurs in never-smokers (2), whereas sqNSCLC is the most common histological type among smokers. However, up to 15% of patients with sqNSCLC have no reported smoking history(3).

Accurate histological classification of NSCLC into major subtypes is critical for treatment selection and prognostic assessment. Despite advancements in diagnostic techniques, misdiagnosis of these subtypes remains a significant challenge in clinical practice. In one study, when compared with preoperative biopsy, only 65.7% of patients undergoing subsequent surgical resection remained classified as having sqNSCLC. Among those with discordant results, approximately 13% (three of 23 re-classified diagnoses) were ultimately diagnosed with LUAD(4). Additionally, a meta-analysis showed that cytological diagnosis of sqNSCLC had a sensitivity of 84% and specificity of 90%, indicating that up to 16% of sqNSCLC diagnoses could be misclassified(5). A study of 93 cases found a 29% disagreement rate among cytopathologists when diagnosing sqNSCLC, further highlighting the subjective nature of histopathological diagnosis(6).

Despite this discordance, recommendations from consensus guidelines frequently rely primarily on histology to determine the utility of molecular testing. Comprehensive genomic profiling (CGP) is an essential component for guiding treatment strategies in patients with advanced NSCLC. The detection of actionable genomic alterations (AGAs) is crucial for determining the most appropriate treatment strategy, which commonly influences the decision between targeted therapy or an immunotherapeutic approach. Furthermore, CMP has potential implications for peri-operative treatment paradigms. Current guidelines recommend that all patients with advanced LUAD undergo CGP with next-generation sequencing (NGS), regardless of clinical characteristics, such as age or smoking history (7). However, the current National Comprehensive Cancer Network (NCCN) guidelines only suggest consideration for molecular testing in sqNSCLC(8). Similarly, the European Society of Medical Oncology (ESMO) clinical practice guidelines recommend against routine CGP in sqNSCLC, except in more unusual cases dictated by age and smoking history (9). As such, routine CGP in sqNSCLC is exceedingly low, with frequent attribution to lack of consensus guideline support(9). Despite this recommendation, numerous AGAs have been observed in lung SCC, including amplifications, fusions, insertions/deletions, and single-nucleotide variants(10,11). Recent advances have expanded the list of targetable genomic alterations in sqNSCLC, which now includes rare but actionable targets, such as ROS1 rearrangements(12,13), RET rearrangements(14,15), and MET alterations(13,16).

Smoking status is known to have a significant impact on the genomic landscape of NSCLC(17). Oncogenic alterations in *EGFR, ALK, ROS1, MET, KRAS, ERBB2*, and *BRAF* have been reported(18,19), and considerable progress has been made in their treatment paradigms(20). However, much of this success has not been entirely recapitulated in sqNSCLC because of the infrequency of events. The frequency of AGAs in non-smokers with sqNSCLC is relatively low but clinically significant(21,22). Various reports have suggested a higher incidence of AGAs in never-smokers with sqNSCLC (3,23,24). In a study conducted by Lindquest et al., NGS analysis revealed that 13% of all patients with sqNSCLC had one potential AGA(25). This, along with the frequency of histologic discordance, suggests a role for broader CGP with NGS in all patients diagnosed with NSCLC, regardless of histology, particularly in non-smokers. We retrospectively assessed the impact of smoking status on the proportion of AGAs in a cohort of sqNSCLC patients treated at a single institution with a descriptive analysis including subsequent clinical outcomes.

## Methods

We retrospectively reviewed liquid biopsy CGP reports from Guardant360 or Guardant360 CDx between October 2020 and July 2023, which were obtained in the context of routine clinical care at a single institution, the City of Hope Comprehensive Cancer Center. Patient data was deidentified and as per City of Hope guidelines with City of Hope Institutional Review Board approval under IRB #23417 and in accordance with the Declaration of Helsinki. Only patients with a final pathologic diagnosis of exclusively sqNSCLC were included in the analysis. Patients with adenosquamous components were excluded. We then obtained social history from clinical documentation. Patients were categorized based on smoking status into the two discrete categories (smoker or non-smoker). AGAs were defined as alterations with FDA-approved therapies for NSCLC. Treatment data, including objective response rate (ORR) as determined by Response Evaluation Criteria in Solid Tumors (RECIST) criteria evaluated by the study team, progression-free survival (PFS) based on radiographic progression of disease with cross-sectional imaging obtained as standard of care and overall survival (OS) based on the date of death as documented by the electronic medical record, were also obtained when available.

## Results

The patient demographics are summarized in Table 1. A total of 56 patients were included in the final analysis. Of these patients, 98% (n = 55) had a documented social history, and 24% (n=13) were classified as non-smokers.

**Table 1.**
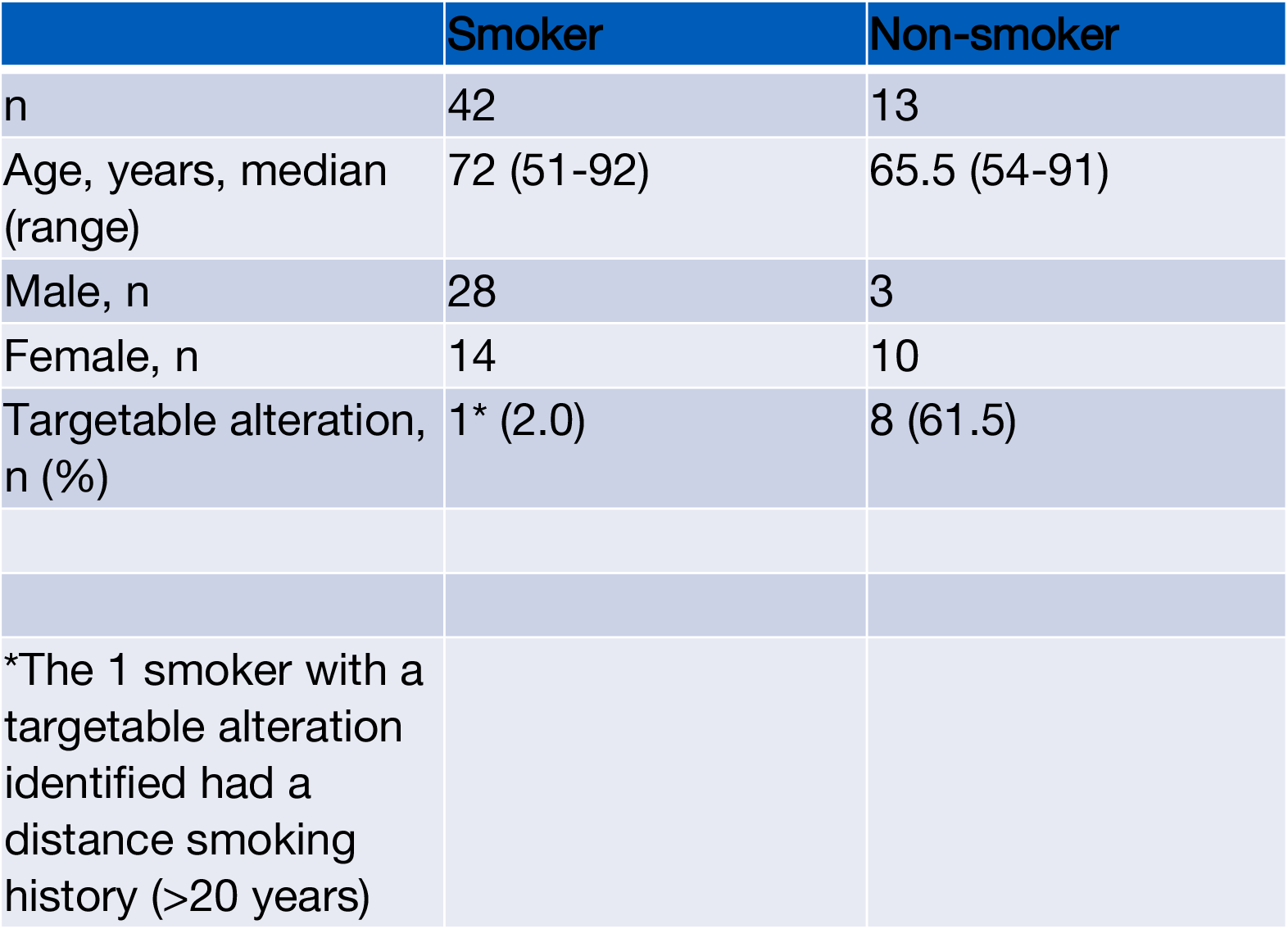

|  | Smoker | Non-smoker |
| --- | --- | --- |
| n | 42 | 13 |
| Age, years, median (range) | 72 (51-92) | 65.5 (54-91) |
| Male, n | 28 | 3 |
| Female, n | 14 | 10 |
| Targetable alteration, n (%) | 1* (2.0) | 8 (61.5) |
| *The 1 smoker with a targetable alteration identified had a distance smoking history (>20 years) |  |  |

In total, 96% (n=54) patients had detectable alterations of variable significance. Of the 13 patients classified as non-smokers, 61.5% (n=8) had an AGA detected by liquid biopsy which had a current FDA-approved therapy. All (100%) patients with AGAs had metastatic disease. Non-smokers were younger and more likely to be female. Of targetable AGAs in the non-smoking cohort, the following alterations were observed: *EGFR* exon 19 deletion (50%, n=4), *MET* exon 14 skipping mutation (25%, n=2), *EGFR* G719S (13%, n=1), *EGFR* E114K (13% n=1). One patient with a former, remote smoking history was also found to have an EGFR exon 19 deletion and who had matched therapy for that alteration (table 2).

**Table 2.**
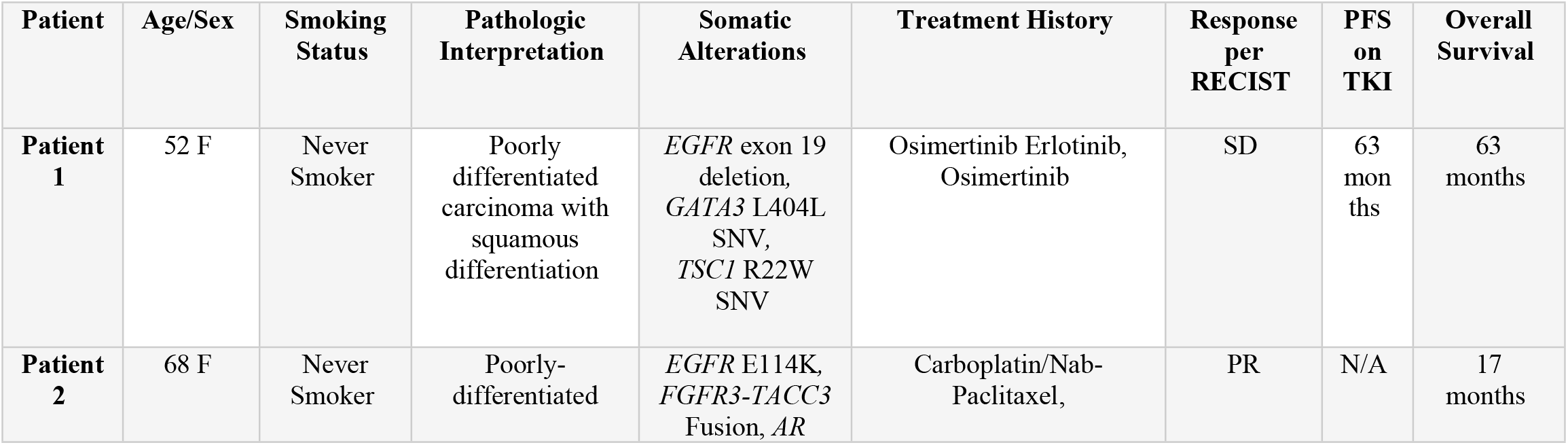
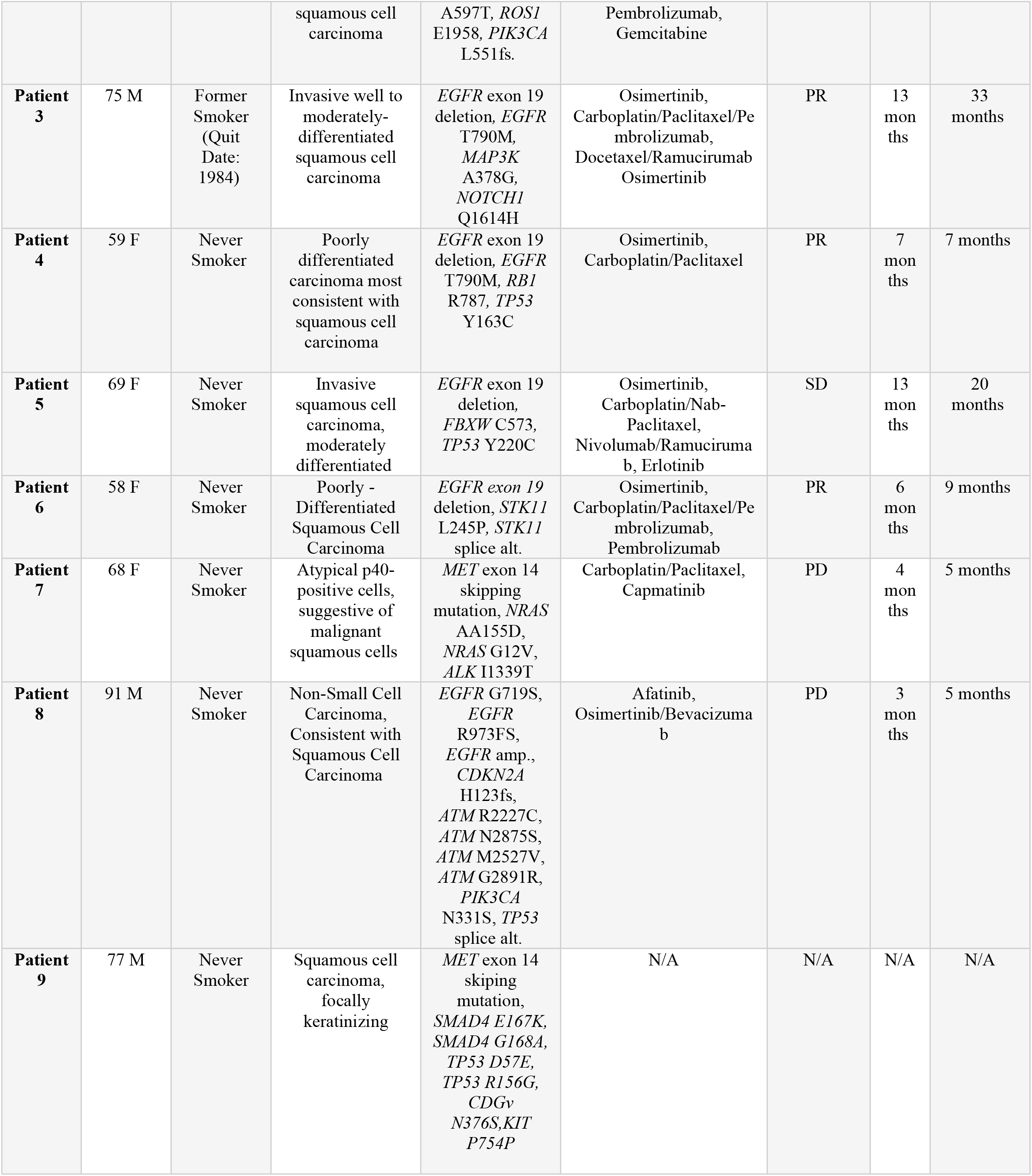
Clinical and social characteristics demonstrated in conjunction with treatment outcomes in patients harboring AGAs.

Among patients with targetable alterations detected, most patients (n=6, 75%) received NCCN guideline-recommended targeted therapy as the first-line treatment. Despite this, the median PFS was 10 months (range: 3-63 months). Osimertinib was the most frequently used targeted therapy, given the proportion of patients with documented *EGFR* alterations. Among those with *EGFR* exon 19 deletions, the median PFS was 13 months (range: 6-63 months).

The social, clinical, and treatment characteristics of all the patients with AGAs are summarized in (Table 2). Most patients with AGA were female (n=6, 75%). This is consistent with *EGFR*-mutant NSCLC, which represents the majority of AGAs observed in our cohort (n=7, 78%). Interestingly, patient 2 had an *EGFR* E114K mutation in the presence of the *FGFR3-TACC3* fusion. This patient did not receive targeted therapy for either atypical *EGFR* alterations or FGFR3-TACC3 fusion. One patient with a *MET* exon 14 skipping mutation died prior to receiving therapy and was excluded from the analysis.

Of the six patients with targetable alterations detected who received matched therapy for that alteration, responses by RECIST criteria as assessed by the clinician included three partial responses (PR), two stable disease (SD), and one progressive disease (PD). The objective response rate (ORR) for this group was 50%, and the clinical benefit rate (CBR) was 83.3%. The clinical and social characteristics of the eight patients are summarized in (Table 2). Most patients with AGAs were female (n=6, 75%).

After excluding Patient 9 with remote smoking history, forty-one patients were classified as current or recent smokers (within 20 years of cessation date). Within this group, the most frequently observed genomic alterations (GA) were *TP53* (n=31, 76%) and *PIK3CA* (n=10, 24%) (Figure 1A). No canonical AGAs affected the *EGFR* gene were observed. In comparison to the non-smoking cohort, there were relatively higher proportions of alterations noted in *TP53* (n=31,76%), *ARID1A* (n=5, 12%), *KRAS* (n=5, 12%), and *KEAP1* (n=4, 10%).

**Figure 1.**
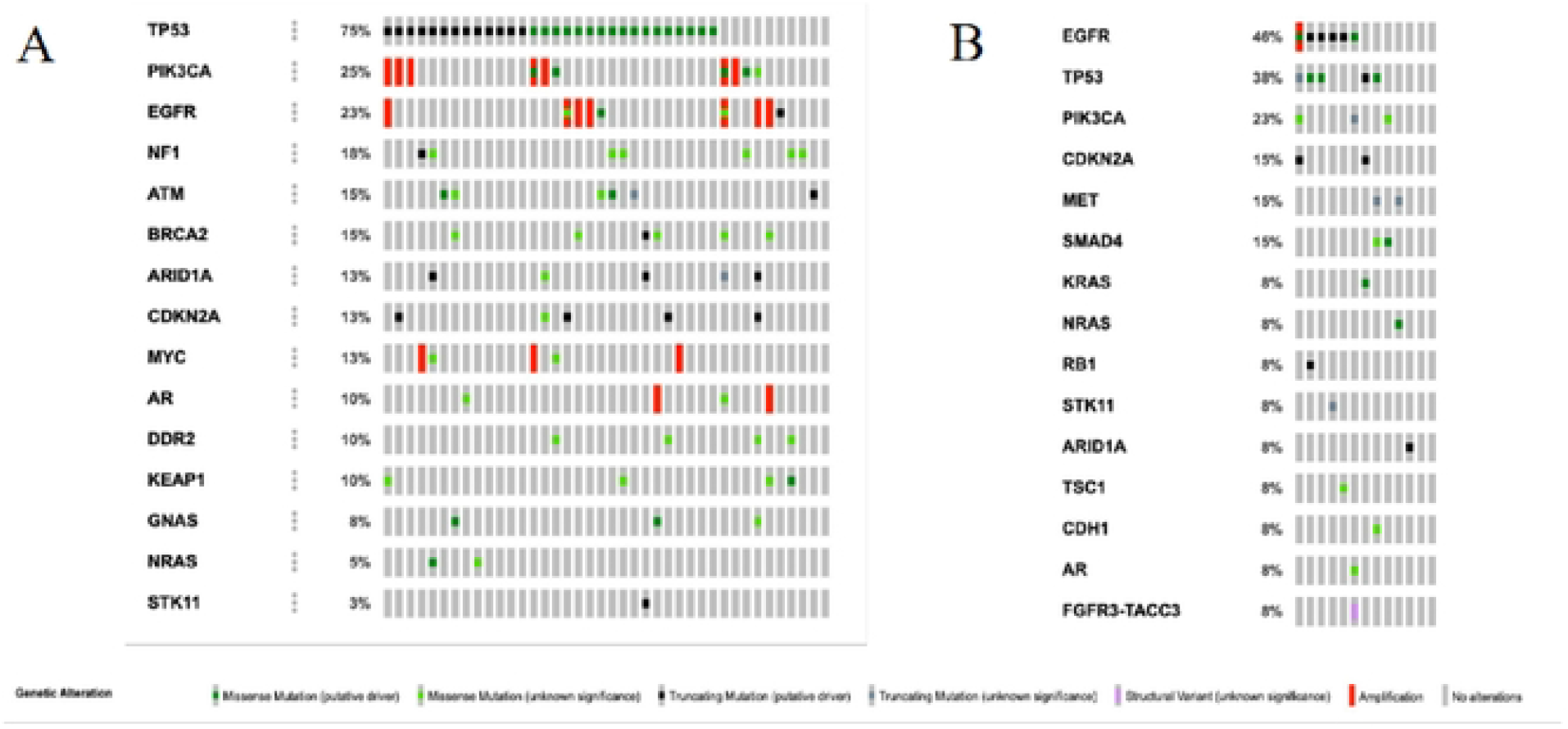
Oncoprint demonstrating genomic alterations in active or recent smokers (A) compared to non-smokers (B).

## Discussion

These data provide valuable insight on the genomic landscape of sqNSCLC based on smoking status and subsequent clinical outcomes following the identification of AGA with CGP. Notably, the majority (61.5%) of non-smokers had AGAs that had a current FDA-approved therapy, with an enrichment of pathogenic *EGFR* and *MET* alterations. Additionally, one patient with a history of remote smoking exposure (quit >20 years ago) was found to have an *EGFR* Exon 19 deletion, suggesting that smoking may need to be considered on a spectrum of chronicity and extent of exposure, as opposed to a binary categorization. In comparison, no patients with a recent or current smoking status were observed to harbor AGAs. These findings are consistent with those of previous studies that have shown the presence of oncogenic alterations in never-smoker sqNSCLC patients (3,26). Additional studies have found that approximately 22% of non-smoker, sqNSCLC patients harbor actionable alteration. Primarily, these were comprised of *EGFR* (7.9%) and *MET* exon 14 skipping (9.5%) alterations with rare *ALK, ROS1, RET* fusions (<2%), which were significantly enriched when compared to their smoking counterparts (<0.5%)(24). This raises the question of whether these are truly exclusively squamous histology with enrichment of AGAs based on smoking status or merely misdiagnosis of an underlying adenosquamous or outright LUAD. Data published by Zhao et al.. al. demonstrated that the prevalence of AGAs, including *EGFR, ALK, MET, RET, ERBB2*, was significantly higher in never-smoker female sqNSCLC patients than that of general sqNSCLC populations evaluated in The Cancer Genome Atlas database (27). Of the non-smokers evaluated in this dataset, 31% of female, non-smoking sqNSCLC patients harbored at least one alteration in *EGFR, ALK, ERBB2*, or *MET*(27).

There are multiple differences in the genomic landscape of smoking-associated versus non-smoking sqNSCLC cited in the literature. Smoking-associated NSCLC exhibit higher tumor mutational burden (TMB) (mean 8.1 mutations/Mb) due to tobacco-induced DNA damage, with dominant C→A transversions (smoking signature) and frequent mutations in *TP53, CDKN2A*, and chromatin modifiers (*DACH1* and *CFTR*)(3,22,28) compared to non-smokers who show lower TMB but enriched for C→T transitions (APOBEC-related) and *EGFR* mutations (7–14% vs. <4% in smokers]). Non-smokers showed higher rates of *EGFR* mutations (up to 70% in some cohorts) and *MET* exon 14 skipping mutations (9.5% vs. 0.4% in smokers)(3,28). Non-smokers also have distinct copy number variations (CNVs) in genes, including *GAB2* amplifications, which are not typically seen in smokers(28). Smoking is postulated to drive TMB, in addition to *KRAS* and *TP53* alterations, whereas non-smoker tumors resemble LUAD genomically. Genomic alterations in smoking-associated sqNSCLC often correlate with reduced immunotherapy efficacy due to higher genomic instability leading to drug resistance and alterations (e.g., *KRAS/STK11* co-mutations) linked to hyperprogressive disease and diminished immune checkpoint inhibitor (ICI) benefits(29). Furthermore, more common alterations in *PIK3CA, ATM*, and MYC are observed at higher frequencies and may negatively affect ICI response (30-32). These considerations have become increasingly relevant with the advent of neoadjuvant chemo-immunotherapeutic approaches in locally advanced NSCLC.

*EGFR*-targeted therapies have become the standard of care in both perioperative and metastatic settings for patients harboring canonical *EGFR* alterations. As expected, clinical trials resulting in FDA approval have mostly been limited to LUAD. In the FLAURA trial, which first demonstrated the superiority of Osimertinib over second-generation *EGFR* TKIs, erlotinib or gefitinib, with an improved PFS (18.9 months vs. 10.2 months) and OS (38.6 months vs. 31.8 months), only 1% of patients were classified as non-adenocarcinoma(33). Similarly, the ADAURA trial, which established the utility of adjuvant osimertinib after platinum-based chemotherapy, had 1% adenosquamous and 2% other histologies (34). More recently, the LAURA trial, consisting of a cohort with only 2% sqNSCLC patients, showed that osimertinib significantly improved progression-free survival by reducing the risk of disease progression or death by 84% compared to placebo (median PFS 39.1 vs. 5.6 months) in patients with unresectable, stage III *EGFR*-mutant NSCLC after definitive chemoradiotherapy(35). In our cohort, all patients with canonical *EGFR* alterations had metastatic disease, with a median PFS of 13 months (range: 6-63 months) observed with osimertinib. A 2017 analysis of TKI therapy in *EGFR*-mutant sqNSCLC demonstrated partial responses in five out of 19 patients (26.3%) and stable disease in four out of 19 patients (21.1%)(36). Furthermore, pilot studies have collectively reported ORRs ranging from 25% to 49% in *EGFR*-mutant sqNSCLC(37). In our cohort, five patients harbored *EGFR* exon 19 deletions, all of whom received targeted therapy with an ORR of 50% and a CBR of 100%.

Analogous to canonical *EGFR* alterations, *EGFR* P-loop and αC-helix compressing (PACC) mutations represent a distinct group of “atypical” alterations resulting in activation of the *EGFR* pathway, resulting in oncogenesis. PACC mutations represent approximately 10-12% of *EGFR-*-mutant NSCLC diagnoses(38,39). Of these, *EGFR* exon 20 insertions, *Gly719Xaa, Leu861Gln*, and *Ser768Ile* represent some of the most frequently occurring and potentially actionable alterations. In a post-hoc analysis of the phase 2 LUX-Lung 2 trial and the phase 3 LUX-Lung 3 and LUX-Lung 6 trials, PACC mutations derived a clinical benefit from the second-generation *EGFR* inhibitor afatinib. Most of this benefit was observed for point mutations rather than *EGFR* exon 20 insertion mutations, with a median PFS of 10.7 months (95% CI 5.6–14·.)(38). Third-generation TKIs, including osimertinib, also demonstrated moderate clinical activity, with prospective data demonstrating an ORR of 50% with a median duration of response of 11.2 months (95% CI, 7.7 to 14.7 months)(40). The activity of both drugs has not been clearly established in sqNSCLC, as prospective trials have specified LUAD as a histological inclusion criterion. In the patients observed to have AGAs in our cohort, two patients (25%) had potentially oncogenic *EGFR* alterations involving atypical alterations including *G719S* and *E114K*. The patient harboring a G719S mutation received front-line targeted therapy with afatinib; however, PD was noted at the first imaging assessment. This patient was also observed to have *EGFR* amplification, which may in part explain the resistance to front-line targeted therapy with afatinib. As noted above, the patient found to have an E114K mutation also harbored an *FGFR3-TACC* fusion with oncogenic potential; however, this patient did not receive targeted therapy for either alteration.

Similarly, *MET* alterations have become a therapeutic target for the development of *MET*-targeting TKIs, capmatinib, tepotinib, and savolitinib. Specifically, *MET* exon 14 skipping mutations show high sensitivity to TKI-based therapy. The GEOMETRY mono-1 trial, which included 8% of sqNSCLC patients, concluded that capmatinib demonstrated meaningful clinical activity and durable responses in *MET* exon 14-altered advanced NSCLC in both front-line and refractory settings (41). A multicenter real-world analysis showed that first-line TKI therapy doubled PFS compared to chemotherapy (11.9 vs. 5.9 months, p<0.002) with a similar improvement in the second-line line (7.7 vs. 4.6 months, p<0.04)(42). In addition, a comparative analysis between GEOMETRY mono-1 trial patients and real-world patients recapitulated these findings demonstrating improved PFS with first-line capmatinib compared with chemotherapy and/or immunotherapy (12.0 vs. 6.2 months, respectively)(43). In our cohort, two patients were observed to harbor a *MET* exon 14 skipping alteration. Unfortunately, one patient died prior to receiving any therapy. Second-line targeted therapy with capmatinib resulted in a PFS of only 4 months, with the first imaging assessment demonstrating PD in the other subject. Potentially, earlier molecular testing and the institution of targeted therapy would have led to better outcomes.

While *FGFR* alterations are frequently observed in NSCLC, potentially actionable *FGFR* fusions comprise approximately 8% of these alterations. *FGFR* fusions have been implicated in oncogenesis in NSCLC, with approximately 1% of these being *FGFR3-TACC* fusions, as observed in Patient 2 within our cohort. Interestingly, these appear to be enriched in sqNSCLC histology(44). Although limited prospective data exist to support *FGFR*-directed therapy, durable responses to erdafitinib in NSCLC have been reported (45). Prospective studies have primarily enrolled patients with *FGFR* amplification, limiting the interpretation of *FGFR* targeted therapies for potentially oncogenic *FGFR* fusions. The SWOG S1400D trial, part of the Lung-MAP protocol, tested the *FGFR* inhibitor AZD4547 in patients with *FGF* pathway-activated sqNSCLC who had progressed after platinum-based therapy. Although AZD4547 was well-tolerated, its clinical efficacy was modest, with a 7% partial response rate, a median PFS of 2.7 months, and an overall survival of 7.5 months(46). Similarly, a phase 1 study of BGJ398, a selective pan-*FGFR* inhibitor, demonstrated only modest activity in *FGFR1*-amplified sqNSCLC, with two patients (12%) achieving PR (two additional patients achieving PR after the data cutoff) and three patients with SD with some degree of tumor regression(47). The RAGNAR study, which evaluated the utility of erdafitinib in *FGFR1-4* mutations or fusions, demonstrated clinically meaningful activity with sqNSCLC, achieving an ORR of 21.4% (95% CI 4.7-50.8)(48).

Although not observed in our cohort, additional AGAs may occur in sqNSCLC that warrant CGP, including *ALK* and *ROS1* rearrangements. The largest study to date evaluating *ALK*-targeting TKIs in *ALK*-rearranged sqNSCLC patients found that these patients obtained meaningful benefit from *ALK*-directed therapy(49). Multiple case reports have documented the responsiveness of *ALK*-fusion sqNSCLC to *ALK*-directed therapies, even after progression on prior chemotherapy(50). Similarly, pure sqNSCLC harboring an oncogenic *EZR-ROS1* fusion has been reported to have a significant and durable CR to crizotinib(51).

The interpretation of these data has significant limitations. Most importantly, pathological diagnosis is frequently established by fine-needle aspiration or core-needle biopsy, which does not effectively exclude the presence of an underlying adenosquamous histology. Furthermore, as all patients with documented AGAs had metastatic disease, the frequency fine needle aspiration (FNA) was high, which could contribute to missed adenosquamous diagnoses. Although most non-smoker patients in our study had AGAs, the sample size was small, which limits the statistical power and may not be representative of the broader sqNSCLC population. Smoking history was obtained from clinical documentation and self-reported data, which may have been subject to recall bias or incomplete reporting. Additionally, as the data were obtained from a single institution, this may limit the generalizability of the findings to broader populations. While response rates to targeted therapy have been reported, this dataset did not allow for in depth interrogation of potential confounding factors that may influence treatment outcome.

In summary, these data support the hypothesis that smoking status can affect the genomic profile of sqNSCLC, potentially having a significant effect on subsequent treatment. Notably, AGAs with FDA-approved therapy were more prevalent in never-smoking patients with sqNSCLC than in smokers, with a significantly higher occurrence of *EGFR* and *MET* alterations. These findings highlight the necessity of incorporating CGP into the initial diagnostic evaluation of newly diagnosed sqNSCLC patients to select appropriate therapeutic interventions. Moreover, the smoking cohort was also observed to have alterations that may impact responsiveness to ICI-based therapies, which is increasingly important with the incorporation of perioperative immunotherapy or chemotherapy-sparing regimens in the metastatic setting. Lastly, given the diagnostic discordance between sqNSCLC, LUAD, and potentially adenosquamous histology, the routine use of CGP appears to be appropriate. Further research is required to assess the clinical impact of smoking history on treatment efficacy in squamous NSCLC and to refine the therapy selection for this distinct patient subgroup.

## Funding Statement

The study was not supported by any grant funding.

## Ethical Compliance

All procedures performed in studies involving human participants were in accordance with the ethical standards of the institutional and/or national research committee and with the 1964 Helsinki Declaration and its later amendments or comparable ethical standards.

## Conflict of Interest declaration

AR reports stock ownership in Merck, Bristol Myers Squibb, honoraria from DAVA Oncology and OMNI Oncology, consulting role in OncoHost. CJ reports stock ownership in Guardant Health. SG, RS, LA, ML, JM, DN, AA, YL, SS, report no conflicts of interest.

## Contribution

AR, SG, CJ were involved in conceptualization, data curation, investigation, methodology, writing – original draft, writing – review & editing. RS, LA, ML, JM, DN, AA, YL, SS were involved in writing – review & editing.

## Notes

**Conflict of Interest Declaration:** AR reports stock ownership in Merck and Bristol Myers Squibb, honoraria from DAVA Oncology and OMNI Oncology, and consulting role in OncoHost. CJ reports stock ownership in Guardant Health. SG, RS, LA, ML, JM, DN, AA, YL, SS report no conflict of interest.

### Competing Interest Statement

I have read the journal's policy and the authors of this manuscript have the following competing interests: AR reports stock ownership in Merck and Bristol Myers Squibb, honoraria from DAVA Oncology and OMNI Oncology, and consulting role in OncoHost. CJ reports stock ownership in Guardant Health. SG, RS, LA, ML, JM, DN, AA, YL, SS report no conflict of interest.

## References

1. Gadgeel SM. New targets in non-small cell lung cancer. Curr Oncol Rep. 2013 Aug;15(4):411–23.

2. Samet JM, Avila-Tang E, Boffetta P, Hannan LM, Olivo-Marston S, Thun MJ, et al. Lung Cancer in Never Smokers: Clinical Epidemiology and Environmental Risk Factors. Clin Cancer Res. 2009 Sep 14;15(18):5626–45.

3. Huang Y, Wang R, Pan Y, Zhang Y, Li H, Cheng C, et al. Clinical and genetic features of lung squamous cell cancer in never-smokers. Oncotarget. 2016 Jun 14;7(24):35979–88.

4. Yamagishi T, Shimizu K, Ochi N, Yamane H, Irei I, Sadahira Y, et al. Histological comparison between preoperative and surgical specimens of non-small cell lung cancer for distinguishing between “squamous” and “non-squamous” cell carcinoma. Diagn Pathol. 2014 May 29;9:103.

5. Layfield LJ, Pearson L, Walker BS, White SK, Schmidt RL. Diagnostic Accuracy of Fine-Needle Aspiration Cytology for Discrimination of Squamous Cell Carcinoma from Adenocarcinoma in Non-Small Cell Lung Cancer: A Systematic Review and Meta-Analysis. Acta Cytol. 2018;62(5–6):318–26.

6. Zakowski MF, Rekhtman N, Auger M, Booth CN, Crothers B, Ghofrani M, et al. Morphologic Accuracy in Differentiating Primary Lung Adenocarcinoma From Squamous Cell Carcinoma in Cytology Specimens. Arch Pathol Lab Med. 2016 Oct;140(10):1116–20.

7. Lindeman NI, Cagle PT, Beasley MB, Chitale DA, Dacic S, Giaccone G, et al. Molecular testing guideline for selection of lung cancer patients for EGFR and ALK tyrosine kinase inhibitors: guideline from the College of American Pathologists, International Association for the Study of Lung Cancer, and Association for Molecular Pathology. J Thorac Oncol Off Publ Int Assoc Study Lung Cancer. 2013 Jul;8(7):823–59.

8. Aisner DL, Riely GJ. Non–Small Cell Lung Cancer: Recommendations for Biomarker Testing and Treatment. J Natl Compr Canc Netw. 2021;19(5.5):610–3.

9. Hendriks LE, Kerr KM, Menis J, Mok TS, Nestle U, Passaro A, et al. Oncogene-addicted metastatic non-small-cell lung cancer: ESMO Clinical Practice Guideline for diagnosis, treatment and follow-up. Ann Oncol Off J Eur Soc Med Oncol. 2023 Apr;34(4):339–57.

10. Lam VK, Tran HT, Banks KC, Lanman RB, Rinsurongkawong W, Peled N, et al. Targeted Tissue and Cell-Free Tumor DNA Sequencing of Advanced Lung Squamous-Cell Carcinoma Reveals Clinically Significant Prevalence of Actionable Alterations. Clin Lung Cancer. 2019;20(1):30–36.e3.

11. Sands JM, Nguyen T, Shivdasani P, Sacher AG, Cheng ML, Alden RS, et al. Next-generation sequencing informs diagnosis and identifies unexpected therapeutic targets in lung squamous cell carcinomas. Lung Cancer. 2020;140:35–41.

12. Gálffy G, Morócz É, Korompay R, Hécz R, Bujdosó R, Puskás R, et al. Targeted therapeutic options in early and metastatic NSCLC-overview. Pathol Oncol Res [Internet]. 2024;30. Available from: https://www.por-journal.com/journals/pathology-and-oncology-research/articles/10.3389/pore.2024.1611715

13. Yang X, Tang Z, Li J, Jiang J, Liu Y. Progress of non-small-cell lung cancer with ROS1 rearrangement. Front Mol Biosci [Internet]. 2023;10. Available from: https://www.frontiersin.org/journals/molecular-biosciences/articles/10.3389/fmolb.2023.1238093

14. Desai A, Menon SP, Dy GK. Alterations in genes other than EGFR/ALK/ROS1 in non-small cell lung cancer: trials and treatment options. Cancer Biol Med. 2016 Mar;13(1):77–86.

15. Gou Q, Gou Q, Gan X, Xie Y. Novel therapeutic strategies for rare mutations in non-small cell lung cancer. Sci Rep. 2024 May 5;14(1):10317.

16. Gendarme S, Bylicki O, Chouaid C, Guisier F. ROS-1 Fusions in Non-Small-Cell Lung Cancer: Evidence to Date. Curr Oncol Tor Ont. 2022 Jan 28;29(2):641–58.

17. Govindan R, Ding L, Griffith M, Subramanian J, Dees ND, Kanchi KL, et al. Genomic landscape of non-small cell lung cancer in smokers and never-smokers. Cell. 2012 Sep 14;150(6):1121–34.

18. Sun Y, Ren Y, Fang Z, Li C, Fang R, Gao B, et al. Lung adenocarcinoma from East Asian never-smokers is a disease largely defined by targetable oncogenic mutant kinases. J Clin Oncol Off J Am Soc Clin Oncol. 2010 Oct 20;28(30):4616–20.

19. Li C, Fang R, Sun Y, Han X, Li F, Gao B, et al. Spectrum of oncogenic driver mutations in lung adenocarcinomas from East Asian never smokers. PloS One. 2011;6(11):e28204.

20. Mok TS, Wu YL, Thongprasert S, Yang CH, Chu DT, Saijo N, et al. Gefitinib or Carboplatin–Paclitaxel in Pulmonary Adenocarcinoma. N Engl J Med. 2009;361(10):947–57.

21. Adib E, Nassar AH, Abou Alaiwi S, Groha S, Akl EW, Sholl LM, et al. Variation in targetable genomic alterations in non-small cell lung cancer by genetic ancestry, sex, smoking history, and histology. Genome Med. 2022 Apr 15;14(1):39.

22. Gou LY, Niu FY, Wu YL, Zhong WZ. Differences in driver genes between smoking-related and non–smoking-related lung cancer in the Chinese population. Cancer. 2015;121(S17):3069–79.

23. Hammerman PS, Lawrence MS, Voet D, Jing R, Cibulskis K, Sivachenko A, et al. Comprehensive genomic characterization of squamous cell lung cancers. Nature. 2012 Sep 1;489(7417):519–25.

24. Reuss JE, Zaemes J, Gandhi N, Walker P, Patel SP, Xiu J, et al. Comprehensive molecular profiling of squamous non-small cell lung cancer reveals high incidence of actionable genomic alterations among patients with no history of smoking. Lung Cancer Amst Neth. 2025 Feb;200:108101.

25. Lindquist KE, Karlsson A, Levéen P, Brunnström H, Reuterswärd C, Holm K, et al. Clinical framework for next generation sequencing based analysis of treatment predictive mutations and multiplexed gene fusion detection in non-small cell lung cancer. Oncotarget. 2017 May 23;8(21):34796–810.

26. Kenmotsu H, Serizawa M, Koh Y, Isaka M, Takahashi T, Taira T, et al. Prospective genetic profiling of squamous cell lung cancer and adenosquamous carcinoma in Japanese patients by multitarget assays. BMC Cancer. 2014 Oct 28;14(1):786.

27. Zhao Y, Dong Y, Zhao R, Zhang B, Wang S, Zhang L, et al. Expression Profiling of Driver Genes in Female Never-smokers With Non-adenocarcinoma Non–small-cell Lung Cancer in China. Clin Lung Cancer. 2020 Sep 1;21(5):e355–62.

28. Park YR, Bae SH, Ji W, Seo EJ, Lee JC, Kim HR, et al. GAB2 Amplification in Squamous Cell Lung Cancer of Non-Smokers. J Korean Med Sci. 2017 Nov;32(11):1784–91.

29. Alessi JV, Elkrief A, Ricciuti B, Wang X, Cortellini A, Vaz VR, et al. Clinicopathologic and Genomic Factors Impacting Efficacy of First-Line Chemoimmunotherapy in Advanced NSCLC. J Thorac Oncol Off Publ Int Assoc Study Lung Cancer. 2023 Jun;18(6):731–43.

30. Chiu M, Lipka MB, Bhateja P, Fu P, Dowlati A. A detailed smoking history and determination of MYC status predict response to checkpoint inhibitors in advanced non-small cell lung cancer. Transl Lung Cancer Res. 2020 Feb;9(1):55–60.

31. Huang YT, Lin X, Liu Y, Chirieac LR, McGovern R, Wain J, et al. Cigarette smoking increases copy number alterations in nonsmall-cell lung cancer. Proc Natl Acad Sci. 2011;108(39):16345–50.

32. Zhou N, Leung CH, William WNJ, Weissferdt A, Pataer A, Godoy MCB, et al. Impact of select actionable genomic alterations on efficacy of neoadjuvant immunotherapy in resectable non-small cell lung cancer. J Immunother Cancer. 2024 Oct 23;12(10).

33. Soria JC, Ohe Y, Vansteenkiste J, Reungwetwattana T, Chewaskulyong B, Lee KH, et al. Osimertinib in Untreated EGFR-Mutated Advanced Non–Small-Cell Lung Cancer. Vol. 378, New England Journal of Medicine. 2018. p. 113–25.

34. Wu YL, Tsuboi M, He J, John T, Grohe C, Majem M, et al. Osimertinib in Resected EGFR-Mutated Non–Small-Cell Lung Cancer. Vol. 383, New England Journal of Medicine. 2020. p. 1711–23.

35. Lu Shun, Kato Terufumi, Dong Xiaorong, Ahn Myung-Ju, Quang Le-Van, Soparattanapaisarn Nopadol, et al. Osimertinib after Chemoradiotherapy in Stage III EGFR-Mutated NSCLC. N Engl J Med. 2024 Aug 14;391(7):585–97.

36. Joshi A, Zanwar S, Noronha V, Patil VM, Chougule A, Kumar R, et al. EGFR mutation in squamous cell carcinoma of the lung: does it carry the same connotation as in adenocarcinomas? OncoTargets Ther. 2017 Mar;Volume 10:1859–63.

37. Jin R, Peng L, Shou J, Wang J, Jin Y, Liang F, et al. EGFR-Mutated Squamous Cell Lung Cancer and Its Association With Outcomes. Front Oncol. 2021;11:680804.

38. Yang JCH, Sequist LV, Geater SL, Tsai CM, Mok TSK, Schuler M, et al. Clinical activity of afatinib in patients with advanced non-small-cell lung cancer harbouring uncommon EGFR mutations: a combined post-hoc analysis of LUX-Lung 2, LUX-Lung 3, and LUX-Lung 6. Lancet Oncol. 2015 Jul;16(7):830–8.

39. Beau-Faller M, Prim N, Ruppert AM, Nanni-Metéllus I, Lacave R, Lacroix L, et al. Rare EGFR exon 18 and exon 20 mutations in non-small-cell lung cancer on 10 117 patients: a multicentre observational study by the French ERMETIC-IFCT network. Ann Oncol Off J Eur Soc Med Oncol. 2014 Jan;25(1):126–31.

40. Cho JH, Lim SH, An HJ, Kim KH, Park KU, Kang EJ, et al. Osimertinib for Patients With Non-Small-Cell Lung Cancer Harboring Uncommon EGFR Mutations: A Multicenter, Open-Label, Phase II Trial (KCSG-LU15-09). J Clin Oncol Off J Am Soc Clin Oncol. 2020 Feb 10;38(5):488–95.

41. Wolf J, Seto T, Han JY, Reguart N, Garon EB, Groen HJM, et al. Capmatinib in MET Exon 14–Mutated or MET-Amplified Non–Small-Cell Lung Cancer. Vol. 383, New England Journal of Medicine. 2020. p. 944–57.

42. Batra U, Singh AK, Nathany S, Dewan A, Sharma M, Amrith BP, et al. Real world experience with MET inhibitors in MET exon 14 skipping mutated non-small cell lung cancer: largest Indian perspective. Discov Oncol. 2025 Mar 8;16(1):286.

43. Socinski MA, Pennell NA, Davies KD. MET Exon 14 Skipping Mutations in Non–Small-Cell Lung Cancer: An Overview of Biology, Clinical Outcomes, and Testing Considerations. JCO Precision Oncology. 2021. p. 653–63.

44. Wang R, Wang L, Li Y, Hu H, Shen L, Shen X, et al. FGFR1/3 Tyrosine Kinase Fusions Define a Unique Molecular Subtype of Non–Small Cell Lung Cancer. Clin Cancer Res. 2014 Jul 31;20(15):4107–14.

45. Pham C, Lang D, Iams WT. Successful Treatment and Retreatment With Erdafitinib for a Patient With FGFR3-TACC3 Fusion Squamous NSCLC: A Case Report. JTO Clin Res Rep. 2023 May 1;4(5):100511.

46. Aggarwal C, Redman MW, Lara PNJ, Borghaei H, Hoffman P, Bradley JD, et al. SWOG S1400D (NCT02965378), a Phase II Study of the Fibroblast Growth Factor Receptor Inhibitor AZD4547 in Previously Treated Patients With Fibroblast Growth Factor Pathway-Activated Stage IV Squamous Cell Lung Cancer (Lung-MAP Substudy). J Thorac Oncol Off Publ Int Assoc Study Lung Cancer. 2019 Oct;14(10):1847–52.

47. Nogova L, Sequist LV, Cassier PA, Hidalgo M, Delord JP, Schuler MH, et al. Targeting FGFR1-amplified lung squamous cell carcinoma with the selective pan-FGFR inhibitor BGJ398. Vol. 32, Journal of Clinical Oncology. 2014. p. 8034–8034.

48. Schuler MH, Tabernero J, Carranza O, Loriot Y, Pant S, Arnold D, et al. Efficacy and safety of erdafitinib in adults with NSCLC and prespecified fibroblast growth factor receptor alterations in the phase 2 open-label, single-arm RAGNAR trial. Vol. 42, Journal of Clinical Oncology. 2024. p. 8515–8515.

49. Wei J, Sun W, Zeng X, Zhou H, Song Z. Efficacy analysis of ALK inhibitors for treating lung squamous carcinoma patients harboring ALK rearrangement. 2024. 2024;16(7):4146–54.

50. Wang W, Song Z, Zhang Y. Response to crizotinib in a squamous cell lung carcinoma patient harbouring echinoderm microtubule-associated protein-like 4-anaplastic lymphoma translocation: A case report. Thorac Cancer. 2016 Apr 26;7(3):355–7.

51. Yakobson A, Mor T, Dina L, Roisman LC, Levin D, Alguayn W, et al. ROS1 in Squamous Non-Small Cell Lung Cancer—Combined Immunotherapy (PD1/CTLA4) or Targeted Therapy? J Cancer Ther. 2020;11(06):365–70.

